# Dynamic gene expression and growth underlie cell-to-cell heterogeneity in *Escherichia coli* stress response

**DOI:** 10.1101/2020.09.14.297101

**Authors:** Nadia M. V. Sampaio, Caroline M. Blassick, Jean-Baptiste Lugagne, Mary J. Dunlop

**Affiliations:** Biomedical Engineering Department, Boston University, Boston, MA 02215; Biological Design Center, Boston University, Boston, MA 02215

## Abstract

Cell-to-cell heterogeneity in gene expression and growth can have critical functional consequences, such as determining whether individual bacteria survive or die following stress. Although phenotypic variability is well documented, the dynamics that underlie it are often unknown. This information is important because dramatically different outcomes can arise from gradual versus rapid changes in expression and growth. Using single-cell time-lapse microscopy, we measured the temporal expression of a suite of stress response reporters in *Escherichia coli*, while simultaneously monitoring growth rate. In conditions without stress, we found several examples of pulsatile expression. Single-cell growth rates were often anti-correlated with gene expression, with changes in growth preceding changes in expression. These expression and growth dynamics have functional consequences, which we demonstrate by measuring survival after challenging cells with the antibiotic ciprofloxacin. Our results suggest that fluctuations in gene expression and growth dynamics in stress response networks can have direct consequences for survival.

## Introduction

Even under otherwise constant environmental conditions, genetically identical cells can display substantial phenotypic heterogeneity, resulting from differences in gene expression and growth rate from cell to cell (*1*–*4*). Phenotypic heterogeneity can, in principle, arise from slow changes within the population where individual cells have different expression levels or growth rates, but maintain their state over many cell cycles. Alternatively, fast dynamics with rapid fluctuations could produce equivalent distributions. The distinction between these alternatives is significant because the timescale over which expression or growth differences persist can ultimately determine if they have functional consequences or are simply short-lived random variations that are filtered before impacting cellular outcomes. Although phenotypic heterogeneity is well documented, the timescales underlying the variation and their ultimate impact on function are often less clear.

We focused on stress response genes in *Escherichia coli* to study the dynamics of expression and growth in single cells. Genes involved in adaptation to stress are among the noisiest genome-wide (*3, 5*). Studies on individual pathways have revealed specific examples in which heterogeneous expression of stress response genes can allow subpopulations of cells to survive sudden environmental stress, such as transient exposure to antimicrobial drugs, oxidative stress, and acid stress (*6*–*9*). Recent studies also indicate that heterogeneity in the expression of genes involved in DNA repair can lead to variability in mutation rates, contributing to microbial evolution (*10*–*12*). Notably, this diversity exists even in the absence of stressors. Although selected studies have begun to reveal examples of how cell-to-cell phenotypic variation can provide important functional capabilities to cell populations, examples of direct links are relatively sparse compared to reports quantifying phenotypic heterogeneity. This motivated our focus on stress response pathways because the effect of the genes involved can be directly assessed by quantifying outcomes like cell survival versus death following stress.

In addition to measurements at a single time point, long-term monitoring of gene expression has begun to uncover examples of rich dynamics in key stress response proteins (*13*). For example, self-cleavage of the regulator LexA produces spontaneous pulses in the SOS response network (*14*). Further, dynamic activity of transcription factors can propagate to downstream genes, with direct consequences for stress tolerance. For instance, pulsatile expression of the transcription factor ComK enables *Bacillus subtilis* cells to enter a transient competent state (*15, 16*). In *E. coli*, heterogeneous expression of RpoS, a key regulator of general stress response, originates from pulses of activation that are inversely correlated with growth, allowing cells to survive oxidative stress (*8*). Additionally, Kim *et al*. demonstrated that genes in the flagellar synthesis network, a process with a pivotal role in microbial pathogenicity, are expressed with different pulsing programs that allow cells to switch between flagellar phenotypes (*17*). Collectively, these studies demonstrate that the temporal dynamics of gene expression play an important role in the regulation of stress response networks.

Growth rate fluctuations have also been observed in single cells across many bacterial species (*2, 8, 18*–*20*). They can arise due to temporal variation in the expression of metabolic enzymes (*2, 21*), the expression of burdensome proteins (*12*), and due to regulatory effects such as feedback involved in cell size control (*19*). These single-cell differences in growth rate can play a functional role in stress tolerance. For example, Narula *et al*. demonstrated that the growth rate of *B. subtilis* under starvation conditions can determine whether individual cells differentiate into spores or remain vegetative (*22*). Single-cell growth rates also play a protective role, for example correlating with differential antibiotic susceptibility in mycobacteria (*23*) and *E. coli* (*7, 24*).

Understanding the timescales associated with gene expression dynamics and growth can provide critical insights into the strategies that cells use to hedge against environmental uncertainty. In this study, we characterize the prevalence and dynamic properties of cell-to-cell phenotypic variation in different branches of the stress response network in *E. coli*. Using time-lapse microscopy, we monitored the activity of key stress response genes, as well as genes involved in biosynthesis and metabolism, in single *E. coli* cells under precisely controlled, unstressed conditions. Our results reveal several new examples of genes that exhibit pulses of gene expression. Furthermore, properties of the dynamics, such as the frequency or amplitude of pulsing, are unique to each gene. Interestingly, fluctuations in the expression of genes frequently occur following variations in growth rate. Finally, focusing on the acid stress regulator GadX, we show that coincident upregulation of *gadX* and reduced growth rate favors tolerance to lethal antibiotic exposure. Together, this work reveals that non-trivial gene expression dynamics are common, even in otherwise constant conditions, and that these dynamic patterns of gene expression can have critical consequences for cell survival.

## Results

### Single-cell measurements of phenotypic heterogeneity

We began by characterizing heterogeneity in the expression of genes with a diverse range of functions (Table S1). These include genes involved in acid resistance (*gadW, gadX, phoP*), multi-drug resistance (*evgA, marA, rob*), heat shock (*rpoH*), oxidative stress response (*oxyR*), SOS response (*dinB, recA, sulA*), and general stress response (*bolA*), in addition to genes involved in biosynthesis and metabolism (*araC, metJ, purA*).

To measure heterogeneity in the expression from each promoter at the single-cell level, we used strains containing transcriptional reporters where the promoter sequence of interest is fused to the coding sequence of a fluorescent protein. We grew independent bulk cultures of each strain to exponential phase and measured heterogeneity across cells in the population using fluorescence microscopy for single cells on agarose pads. Cell-to-cell differences in gene expression resulted in different distributions of reporter levels for each gene (Fig. 1A, Fig. S1). Measurements of the reporters revealed many instances of wide distributions, indicative of a broad range of expression levels across cells within a population. Distributions tended to skew to the right of the mean (skew > 0) due to the presence of highly-expressing cells.

**Figure 1.**
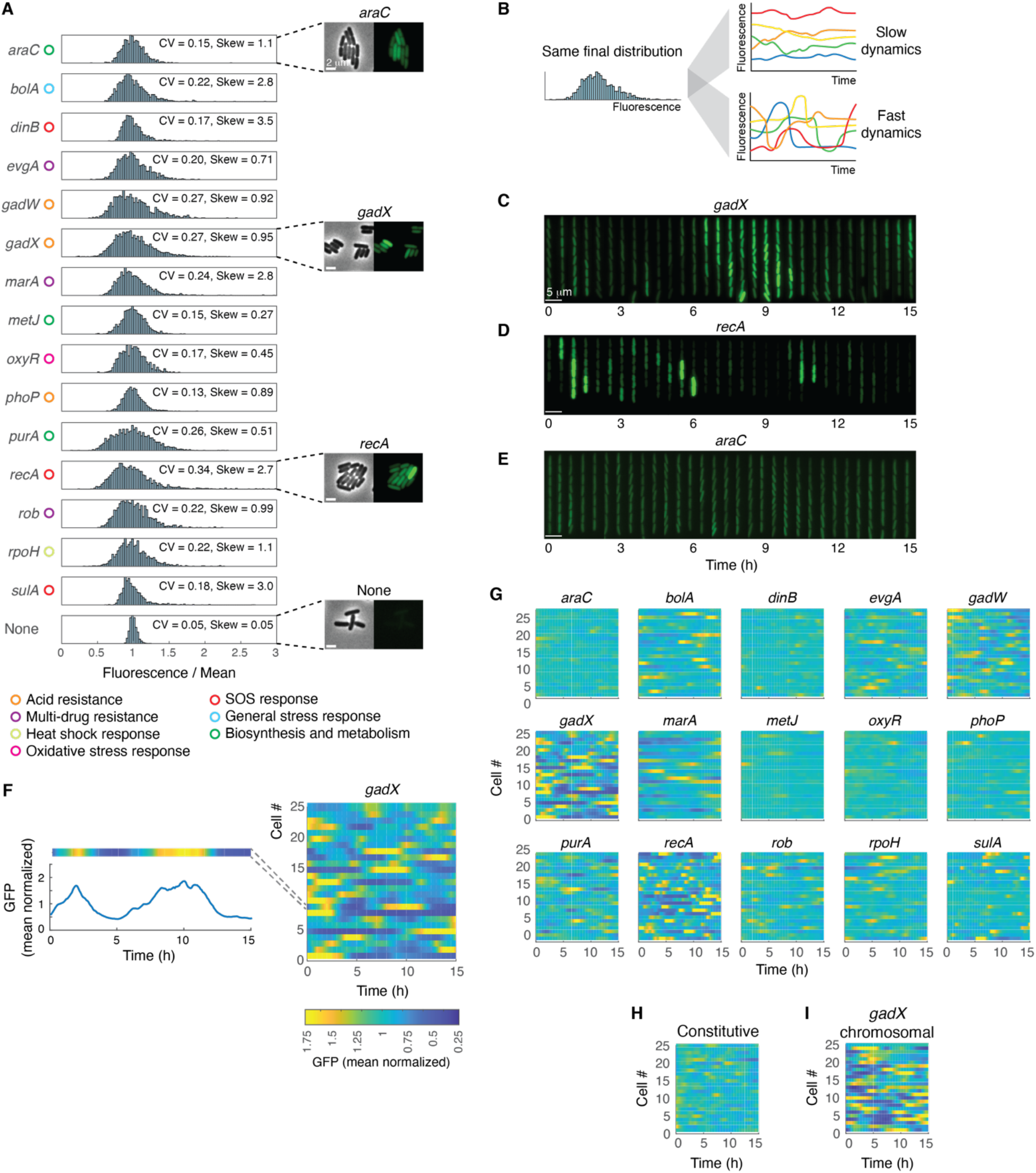
Cell-to-cell heterogeneity and temporal dynamics of gene expression. **(A)** Histograms of fluorescence values show cell-to-cell variation. Values are presented normalized relative to their means to allow for comparison across reporters. The same data without normalization are shown in Fig. S1. Corresponding coefficient of variation (CV) values obtained from snapshot images of cells grown in bulk cultures are listed in the figure. The skewness (skew) of the distribution, which is positive for distributions that are right-skewed, is also listed. Insets show phase contrast and fluorescence images for representative strains. Functional classes are listed for each gene. Scale bar, 2 μm. **(B)** Schematic representation of fluorescence signal over time originating from single cells with slow or fast dynamics of gene expression that result in identical fluorescence distributions for the final timepoint. **(C-E)** Representative kymographs of cells containing transcriptional fluorescent reporters for **(C)** *gadX*, **(D)** *recA*, and **(E)** *araC*. In all cases, fluorescence values are normalized to the mean to allow comparisons across the reporters. Scale bar, 5 μm. **(F)** Single-cell measurements of green fluorescent protein expression (GFP) over time for the *gadX* reporter. Colored heat maps summarize the time-series data. Data from 25 cells are shown, however this represents a subset of the time-series data. **(G)** Heat maps summarizing the temporal dynamics of gene expression for all reporters. **(H)** Constitutive reporter heat map. **(I)** Chromosomally-integrated *gadX* reporter heat map. Color scale and normalization approach in (F) applies to all heat map data.

However, measures of static distributions do not reveal the underlying dynamics that generate them, prompting the question: Do these distributions stem from long-lived fixed subpopulations with slow dynamics or from conditions with fast dynamics where individual cells transition between different expression levels over time (Fig. 1B)? The significance of this question is particularly pertinent in the context of stress response networks, where the timescales over which individual genes are active may have concrete implications for tolerance levels. For instance, if a transcription factor that activates genes involved in stress response exhibits short pulses in expression, these pulses might be insufficient to turn on expression of downstream genes, while more sustained expression could. Further, single-cell growth rates can impact survival, thus the interplay between expression and growth may be significant for determining tolerance.

### Pulsatile gene expression can underlie cell-to-cell heterogeneity

Thus, we next aimed to quantify the temporal dynamics underlying the distributions of gene expression and growth. We measured expression in cell lineages over many generations using a ‘mother machine’ microfluidic device (*25*) and time-lapse fluorescence microscopy. In this device, ‘mother’ cells are trapped at the top of one-ended chambers and maintained indefinitely in exponential phase through the addition of fresh media, allowing for multi-hour imaging of cell lineages. We used this to monitor gene expression in tens to hundreds of independent cell lineages for each reporter for at least 15 hours. To quantify gene expression over time, we used our deep learning based cell segmentation and tracking algorithm (*26*) to extract single-cell resolution data from the mother cell and its progeny.

We observed heterogeneity in gene expression for cells growing in these precisely controlled conditions, which was consistent with measurements acquired from bulk culture snapshots (Fig. S2). Long-term monitoring of expression revealed a diversity of phenotypes, with some genes that fluctuated at a low level, likely due to variation in copy number in the reporter plasmid and inherent stochasticity in gene expression, while others were highly dynamic in their expression (Fig. 1C-E). We observed that variability within a distribution often arose from pulsatile dynamics, with single cells transitioning smoothly between different expression states. These results join other recent single-cell studies showing transcriptional pulses in expression in the absence of stress (*8, 14, 17*). We found that the timescales of these fluctuations were specific to each gene, and the pulses themselves ranged in intensity. For instance, *gadX*, which encodes a transcription factor that regulates ∼34 genes in the acid resistance system, exhibits pulses of high amplitude and duration that persist well beyond the cell cycle length, as visible in multi-generation patterns of gene expression (Fig. 1C, Movie S1). In contrast, *recA*, which plays a central role in the processes of homologous recombination and SOS response, showed large amplitude pulses, but with shorter durations (Fig. 1D, Movie S2). Yet others, like *araC*, which encodes a transcription factor that regulates arabinose catabolism and transport, were more muted in their changes, and exhibited only mild, low amplitude fluctuations (Fig. 1E, Movie S3). Surveying 15 reporters, we observed a wide range of temporal gene expression profiles (Fig. 1F-G). In all cases, the mean fluorescence levels were consistent across the duration of the experiments despite the fluctuations in gene expression at the single-cell level (Fig. S3). Importantly, the dynamic activity observed for many reporters occurred under constant growth conditions, where parameters such as pH, temperature, and growth medium do not fluctuate.

As a control, we also included a plasmid-based constitutive reporter to quantify the baseline level of fluctuations expected from variation in plasmid copy number (Fig. 1H). The constitutive reporter exhibits mild changes in expression but does not reach the level of variation we observed in reporters like *gadX* and *recA*. As an additional control, we also chromosomally integrated the *gadX* reporter and confirmed that pulses in gene expression in cells containing the plasmid and chromosomally-integrated versions of the reporters were comparable (Fig. 1I). Low expression levels prevented us from conducting all experiments using chromosomal reporters, however the constitutive reporter serves as a control to establish baseline effects that are plasmid-based rather than specific to the stress response reporters.

### Properties of temporal expression vary across the reporters

Next, we defined several metrics across which to assess dynamic behavior. We selected properties related to pulses including their frequency, duration, and amplitude (Fig. 2A). Pulse frequencies were broadly distributed, ranging from examples that exceed 0.25 pulses per hour (one pulse every four hours) for *recA* to much less frequent conditions where our peak-finding algorithm rarely identified pulses (Fig. 2B). Pulse durations typically ranged from 0 - 4 hours, and showed instances of precision (e.g. *recA, sulA*) in addition to examples with widely variable durations (e.g. *bolA, evgA, rpoH*). Pulse amplitudes were predominantly small, with average amplitudes around 75% of the mean, or 0.75x, but there were notable instances where the distributions had long tails such that amplitudes extended well above this for a subset of the pulses. For example, *gadX* and *recA* exhibit pulses that significantly deviate from the mean. To alleviate the potential concern that these relationships were a result of the specific method by which we identified pulses, we repeated the calculations under a range of different peak determination thresholds and found that our results were not sensitive to the precise threshold definition, provided the threshold was set high enough to exclude most fluctuations in the constitutive reporter control (Fig. S4).

**Figure 2.**
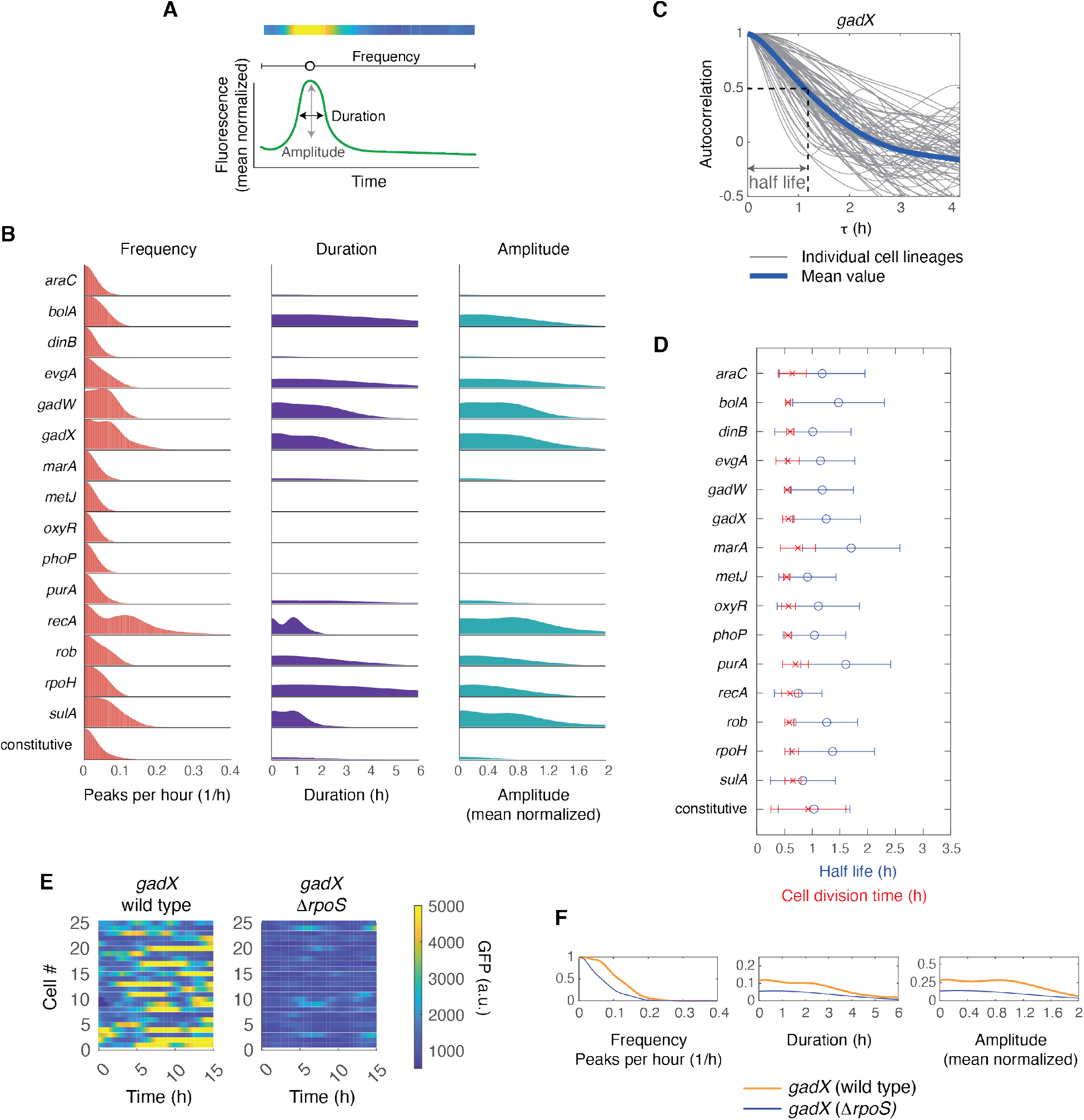
Pulsing dynamics vary across the reporters. **(A)** Schematic representation of pulse characteristics. **(B)** Distributions of pulse frequency, duration, and amplitude for all reporters. Note that typical movie durations are 15 - 20 hours (Table S2), thus we are limited in our ability to measure very low frequencies accurately (1/15 = 0.067). Duration and amplitude histograms are normalized such that the maximal height corresponds to the fraction of lineages with at least one pulse. **(C)** Autocorrelation of the fluorescence signal for independent cell lineages with the *gadX* reporter. Half life is defined as the value of the time shift τ when the mean of the autocorrelation curve crosses 0.5. **(D)** Cell division time and half life for each of the reporters. Error bars show standard deviation around the mean. **(E)** Heat maps summarizing single-cell measurements of green fluorescent protein expression (GFP) over time for the *gadX* reporter in wild type and Δ*rpoS* cells. Identical reporters are used in both strains, thus the unnormalized data can be compared directly. Normalized versions are shown in Fig. S7. **(F)** Frequency, duration, and amplitude distributions for the *gadX* reporter in the wild type and Δ*rpoS* backgrounds.

We next asked what characteristic timescales the gene expression dynamics exhibited by calculating the autocorrelation of the fluorescence signal. In all cases, we observed monotonically decreasing autocorrelation curves, ruling out the presence of regular period signals, such as oscillatory behavior (Fig. S5). We calculated the half life associated with the autocorrelation curve for each gene (Fig. 2C-D). If the fluorescent reporter levels decrease solely due to dilution resulting from growth and division, the half life will equal the cell division time. Longer half lives can indicate the presence of memory, for example due to regulatory networks that causes signals to persist. We found that half lives were always greater than or equal to the cell division time, consistent with the use of stable fluorescent reporters (Fig. 2D). Notably, we observed cases where the average exceeded the cell division time by 2-3-fold, potentially indicative of memory within the network. We verified that these calculations performed on data from mother cells were consistent with correlations within the lineage tree. Indeed, tracking fluorescence signals from mother to daughter to granddaughter cells revealed a strong positive correlation that persisted across multiple generations (Fig. S6).

### RpoS levels influence *gadX* pulsing dynamics

One question that our results provoke is why some of the reporters exhibit pulsatile expression. We examined this in more detail for the case of the *gadX* reporter. RpoS plays a role in *gadX* regulation and has been shown to exhibit pulsatile dynamics during exponential growth (*8, 27, 28*). Thus, we hypothesized that fluctuations in cellular RpoS levels during exponential growth could be underlying *gadX* pulsing. We used the same *gadX* reporter and compared expression in wild type and Δ*rpoS* strains. Deletion of *rpoS* reduced *gadX* expression and fluctuations in *gadX* expression became significantly less noticeable (Fig. 2E, Fig. S7). The frequency, amplitude, and duration of *gadX* pulses in the Δ*rpoS* cells were reduced relative to the characteristic profile of wild type cells (Fig. 2F), indicating a direct impact of RpoS levels on *gadX* pulsing dynamics. Fluctuations in the availability of the master regulator RpoS may also contribute to the profiles observed for other downstream genes that RpoS controls. The mechanism leading to pulsing dynamics of other genes involved the SOS response, like *recA* and *sulA*, has been shown to originate from fluctuations in the availability of their common negative regulator LexA (*14*). Thus, fluctuations in master regulators of gene expression might be a common mechanism leading to heterogeneous expression of downstream genes.

### Pulsatile expression dynamics and growth

We next asked whether growth rate contributes to the observed variability in expression levels. While the growth rate of individual *E. coli* cells shows long-term stability during replicative aging (*25*), it can exhibit noisy temporal behavior (*2, 8, 19*). Patange *et al*. demonstrated that growth rates fluctuate, with pulses of slow growth that last for several generations and are anti-correlated with pulses of activation of RpoS (*8*). However, it is not clear whether this feature is specific to RpoS or if it is common to many genes. We asked whether the pulsatile dynamics we observed coincided with dynamic patterns in growth rate. To test this, we extracted the instantaneous growth rate from cell lineages over time and compared them to fluorescence data for the different reporters.

In principle, fluorescence levels, which report underlying cellular properties such as those related to metabolism, can precede changes in growth. Alternatively, growth can drive changes in fluorescence since it affects dilution rates; regulatory links between growth rate and fluorescence are also possible (Fig. 3A). In both cases, these relationships could be positive or negative depending on the precise underlying mechanism.

**Figure 3.**
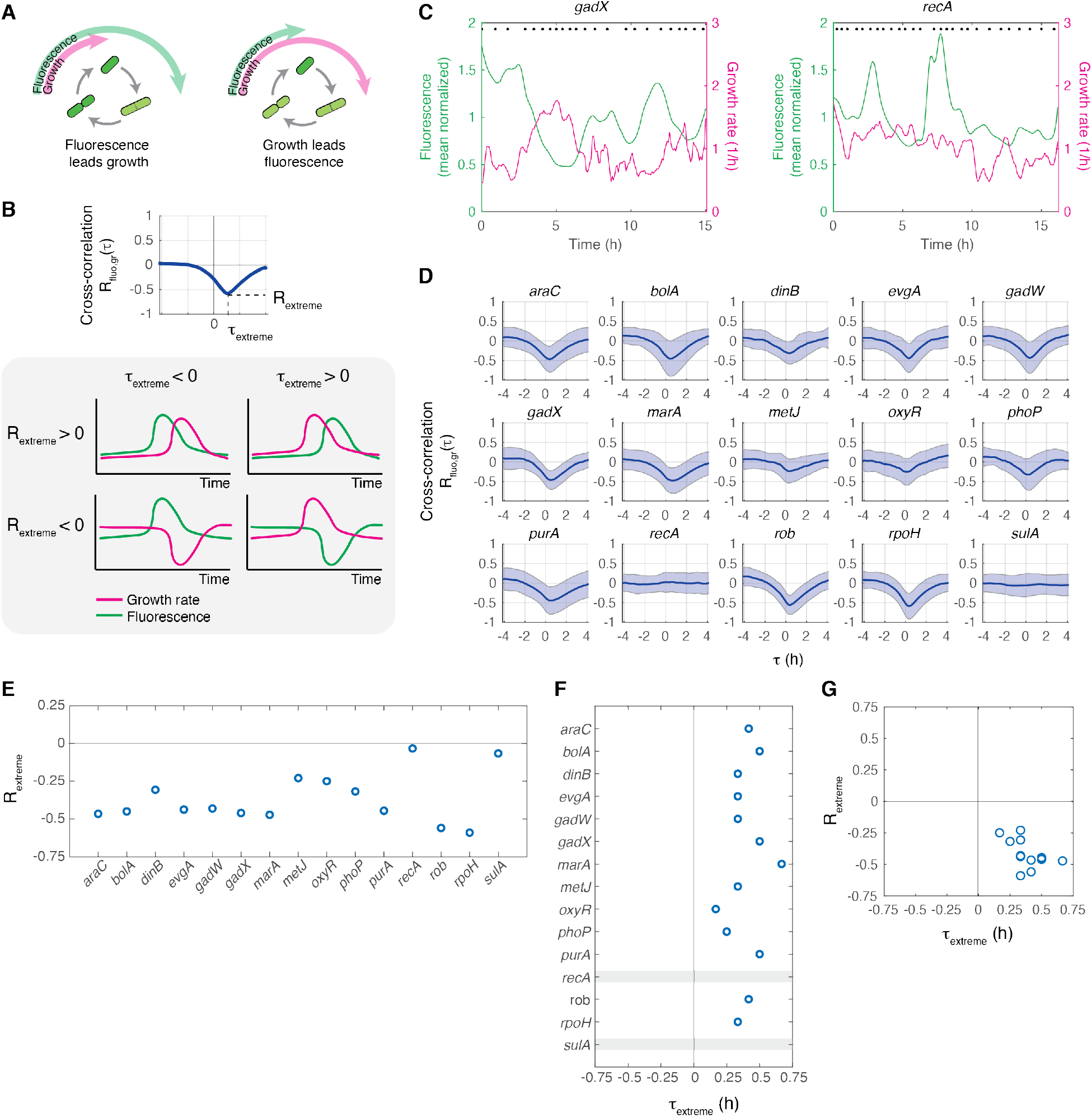
Growth and gene expression dynamics are often anti-correlated. **(A)** Two possible models for the relationship between fluorescent reporter levels and growth. **(B)** Schematic of cross-correlation function R_fluo,gr_(τ) indicating points corresponding to R_extreme_ and τ_extreme_ (top). Temporal patterns of the fluorescence and growth rate and their impact on R_extreme_ and τ_extreme_ (bottom). **(C)** Representative fluorescence (green) and growth rate (magenta) values over time for a single mother cell with a reporter for *gadX* (left) or *recA* (right). Black dots at the top of the figure indicate cell division events. Data are smoothed with a moving average filter with a window of 5 frames. **(D)** Cross-correlations for all reporters. Shaded region represents standard deviation about the mean. **(E)** R_extreme_ and **(F)** τ_extreme_ values for each reporter. **(G)** R_extreme_ as a function of τ_extreme_. For signals without a peak (|R_extreme_| < 0.1) we do not calculate a τ_extreme_ value.

To distinguish between these possibilities, we computed temporal cross-correlations between fluorescence and growth rate, R_fluo,gr_(τ) (Fig. 3B). The cross-correlation measures how well the fluorescence and growth signals are correlated when the growth signal is shifted by a time τ relative to the fluorescence signal. Cross-correlation curves for our reporters have characteristic shapes, where signals are uncorrelated at large positive and negative values of the time shift, τ. For intermediate values of τ we regularly observed valleys in the cross-correlation. We defined R_extreme_ as the point on the cross-correlation with the largest absolute value, indicating the largest correlation between the fluorescence and growth rate signals. R_extreme_ can be either positive or negative, depending on whether the signals are correlated or anti-correlated. We also defined τ_extreme_ as the time shift associated with R_extreme_, where τ_extreme_ < 0 when changes in fluorescence precede changes in growth and τ_extreme_ > 0 when growth leads fluorescence.

### Growth and expression are often anti-correlated with growth leading expression

We found examples where a strong anti-correlation between fluorescence and growth were visible, such as with the *gadX* reporter, and other cases where the signals were uncorrelated, such as with the *recA* reporter (Fig. 3C). To quantify this trend across many cell lineages, we calculated cross-correlations between growth and fluorescence for all reporters (Fig. 3D). In many cases we observed an anti-correlation with a positive time shift between growth rate and gene expression, indicating that the pulses of expression from these promoters were preceded by a decrease in growth rate. In addition, we confirmed that the chromosomally-integrated version of the *gadX* reporter produces cross-correlation curves that are similar to the plasmid-based version (Fig. S8 A-B). Although the accumulation of fluorescent proteins resulting from reduced dilution due to cell growth and division could potentially artificially generate this relationship, we observed only a modest anti-correlation for several of the promoters, a near zero or mildly positive relationship for the constitutive reporter (Fig. S8 C-D), and no anti-correlation for *recA* and *sulA* (Fig. 3E). Thus, it is unlikely that these results are due solely to a reduced dilution rate.

Changes in growth rate never lagged changes in fluorescence for all reporters we measured (τ_extreme_ > 0) (Fig. 3F). It is possible that fluorophore maturation times could systematically introduce a lag between actual promoter activity and read-out of the fluorescent protein. Maturation times for our reporters are relatively rapid, with 50% of fluorescent proteins maturing within ∼6 minutes (*29*), however reporter maturation times could systematically introduce a small positive shift in τ_extreme_. Overall, our measurements indicate that even cells growing exponentially under optimal conditions undergo episodes of slow growth that are largely followed by the pulses of stress-related genes, which frequently demonstrate an inverse relationship between growth rate and fluorescence (Fig 3G).

We also measured correlations between fluorescence and cell length for all the reporters (Fig. S9). These signals were not strongly correlated for most of the reporters. However, we did observe positive correlations between expression of *recA* and *sulA* and cell length. This result is expected, because these genes play a key role in the SOS response, which is known to induce cell filamentation (*30*–*32*). Even in unstressed conditions, we find that a small subset of cells exist in a state with high gene expression and corresponding elongated cell morphology.

### Gene expression and growth dynamics transiently protect cells against stress

A critical question is whether these changes in gene expression and growth have concrete implications for whether a single cell will survive or die following stress. In other words, are the dynamics we observed sufficient to provide meaningful phenotypic differences that cause a cell to tolerate or succumb to stress? Slow growth and induction of stress response genes have previously been reported as mechanisms that allow survival at the populational level (*33*–*37*). This prompted us to investigate whether pulses in gene expression and growth influence the chances of survival under stress. Further, we asked whether survival was the result of a fortuitous condition of gene expression or slow growth upon stress introduction, or if a cell’s past history was important.

For these studies, we focused on expression of *gadX*, as our measurements demonstrate that this reporter is expressed in large pulses and has a strong anti-correlation with growth. We previously demonstrated that *gadX* is heterogeneously expressed in the absence of antibiotic stress and that expression levels correlate with longer survival times under constant carbenicillin exposure (*7*). However, it was not clear how prior history or pulsing dynamics contribute to this phenotype and whether cells could recover normal cell division after transient drug treatment. Here, we quantified how *gadX* fluctuations preceding sudden exposure to a lethal antibiotic dose influence survival. To do so, we leveraged the mother machine device to rapidly switch input media and exposed exponentially dividing cells to a short pulse of ciprofloxacin, a fluoroquinolone drug widely used to treat bacterial infections. We treated cells with 2 μg/ml ciprofloxacin, which corresponds to 100x the minimum inhibitory concentration (MIC), for 35 minutes before switching back to growth medium without antibiotics. This experimental set up allowed us to monitor the dynamics of *gadX* expression and growth in single cells prior to antibiotic treatment while also recording the outcome of each cell lineage.

Within the same experiment, we observed instances where single cells were able to survive ciprofloxacin treatment (Fig. 4A, Movie S4) and cases where treatment killed the cells (Fig. 4B, Movie S5). In addition to surviving and dying, we also observed a third category of outcomes where cells filamented. However, it was difficult to accurately assess whether these cells survived or died, as they were frequently swept out of the mother machine chamber and lost from the field of view. Thus, we focused our analysis on the surviving and dying cells because we could accurately determine their outcomes. By tracking gene expression history preceding antibiotic treatment, we observed pulses in fluorescence from the *gadX* reporter prior to ciprofloxacin addition in both the surviving and dying cells (Fig. 4C). Cells that exhibited expression pulses in the past, but had low expression at the time of ciprofloxacin treatment were more likely to die. In contrast, cells which were in the fortuitous state of having an ongoing *gadX* pulse at the time of ciprofloxacin addition were more likely to exhibit transient stress tolerance.

**Figure 4.**
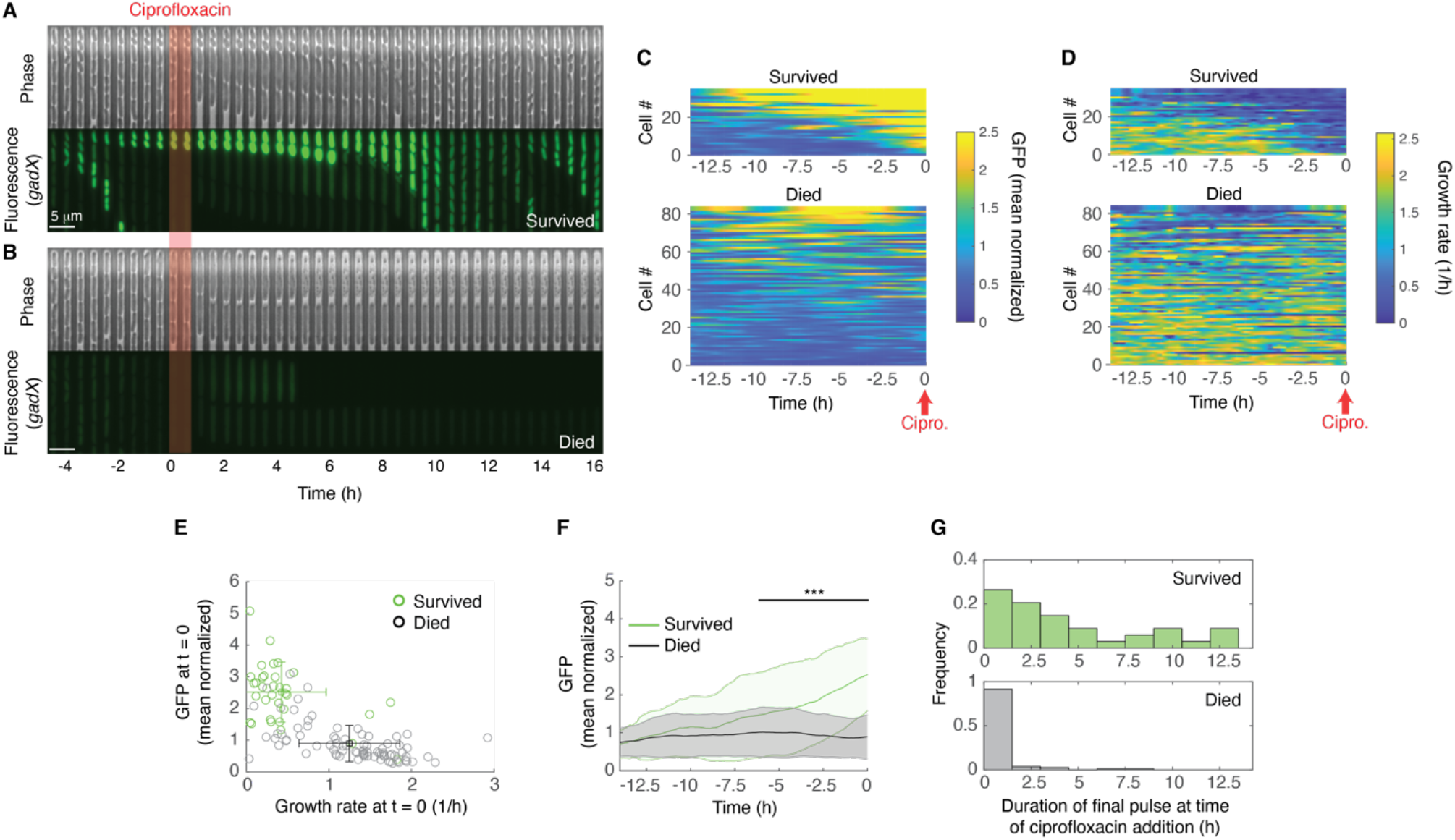
Cell-to-cell differences in *gadX* expression and growth influence survival outcomes after antibiotic exposure. **(A)** Representative kymograph of a cell lineage that survives and **(B)** dies after antibiotic treatment. Images are from nearby chambers within the same microfluidic chip. Shaded area in red indicates 35 minute period of ciprofloxacin treatment. **(C)** Single-cell *gadX* expression and **(D)** growth rate for the period preceding ciprofloxacin treatment. Cell numbers in (C) and (D) correspond to the same cells. Time-series are sorted by the mean value of the fluorescence signal for t < 0, from high to low. **(E)** Individual cells that survived (green) or died (gray) following ciprofloxacin treatment plotted as a function of growth rate and fluorescence at t = 0. Error bars show standard deviation about the mean. **(F)** Mean fluorescence from the *gadX* reporter for the period preceding ciprofloxacin addition. Shaded regions correspond to standard deviation about the mean. Statistics are conducted at each timepoint; the horizontal bar indicates all time points where *** P < 0.001, two-sample *t* test for differences between the mean of the cells that survived and died. **(G)** Histogram of duration of the pulse that cells were in upon ciprofloxacin addition, sorted by outcome.

We also looked at cellular outcome as a function of the growth rate. Consistent with our results showing anti-correlations between fluorescence and growth for *gadX* (Fig. 3D), we observed decreased growth rates in cells that survived, which were associated with increases in fluorescence signals (Fig. 4D). However, reduced growth rate alone does not appear to be sufficient for survival, as we observed cells that died which had similar growth rates to those that survived. Measurements of the instantaneous fluorescence and growth rate at t = 0, just prior to antibiotic addition, show that most surviving cells exhibited both elevated *gadX* expression and reduced growth rate immediately prior to ciprofloxacin treatment (Fig. 4E).

By looking at gene expression history prior to ciprofloxacin addition, we see clear differences emerging between the populations of cells that will survive and die ∼6 hours prior to treatment (Fig. 4F). The trend we observe does not reflect a linear increase in fluorescence in individual cells, but rather results from *gadX* pulsing dynamics. For surviving and dying cohorts, we quantified how long each cell had been experiencing its final pulse of *gadX* expression at the point when ciprofloxacin was introduced (Fig. 4G). We found that cells that survived were far more likely to be in the midst of a *gadX* expression pulse than those that died. Pulse duration did not have a strong impact on survival; simply being within a pulse was often sufficient for survival.

Neither growth rate nor *gadX* expression alone is expected to completely determine cellular outcome, as stress response is a complex process; instead, our results suggest that pulsing dynamics can bias the probability that a single cell will live or die. Cell-to-cell heterogeneity in *gadX* expression is the result of pulsing dynamics, which produce diverse distributions of cells. Thus, we demonstrate that *gadX* pulsing can play a protective role against ciprofloxacin stress, indicating a clear link between single-cell dynamics and cell fate.

## Discussion

Although cells can display substantial phenotypic heterogeneity in constant environmental conditions, the timescales over which gene expression levels and growth rate fluctuate and how this ultimately impacts function are often less clear. To provide insight into these questions, we focused on genes involved in stress response in *E. coli*. Our results demonstrate that distributions of gene expression levels and growth rates can originate from rich dynamic activity, where individual cells transition between expression levels over time. Our findings expand previously published observations on pulsing dynamics (*8, 13, 14, 17, 38*) to a broader set of genes, suggesting that these time-varying differences in expression and growth may be more common than previously appreciated. These fluctuations occur under uniform, unstressed conditions and are observed across genes with diverse stress response roles. Notably, temporal changes in gene expression are also related to growth. In our measurements, we found many instances where pulses of high activity followed decreases in growth rate. However, this link is not universal, which indicates that this effect is not an unavoidable consequence of growth, and rather that the expression and growth relationship is specific to each gene. Finally, we asked whether gene expression dynamics have functional consequences, focusing on expression of *gadX*. We showed that a functional outcome of *gadX* pulsing and the corresponding decrease in growth rate is that cells are more likely to survive following sudden exposure to a lethal dose of ciprofloxacin.

Our results with *gadX* join a small number of other studies demonstrating examples of functional phenotypic heterogeneity. For example, the stochastic activation of diverse stress response genes, such as those encoding porins (*39*), the catalase-peroxidase KatG (*40*), and the multiple antibiotic resistance activator MarA (*6, 41, 42*) result in transient tolerance to exogenous stress. Other examples include biased partitioning of efflux pumps during cell division that results in differential antibiotic susceptibility (*43*) and heterogeneous induction of *gad* regulon genes during antibiotic treatment that cross-protects cells against subsequent acid stress (*9*).

Reduced growth rates can also enable cells to transiently resist stress, for example, by playing a major role in the formation of persister cells (*34, 44*). By tracking expression dynamics and growth rates simultaneously, we found many instances in which these metrics are inversely correlated over time. Interestingly, an analogous observation was reported by Patange *et al*. (*8*) that demonstrated that the stress response master regulator RpoS is expressed with pulsatile dynamics in exponentially dividing cells and is inversely correlated with growth rate. We demonstrated that RpoS availability directly contributes to *gadX* pulsing dynamics, and could potentially serve as a common mechanism underlying the expression and growth dynamics associated with many additional genes characterized in our study. Other mechanisms are also possible, as some genes in our study with strong growth rate anti-correlations (e.g. *marA*) are not known to be regulated by RpoS. In some instances other mechanisms have been demonstrated, as in a recent study that indicated that self-degradation of the SOS response repressor LexA triggers pulses in the expression of *recA* and *sulA* during exponential growth in unperturbed conditions (*14*). Fluctuations in LexA result in frequent pulses in *recA* expression, while *sulA* is less susceptible to LexA variability, corroborating our observations. It is also noteworthy that, in addition to RpoS, at least one additional sigma factor, RpoH, was activated with pulsatile dynamics during exponential growth, a behavior that can potentially be propagated to >150 downstream genes that collectively control the cellular heat-shock response (*45, 46*). The pulsatile activity of these regulators might indicate that the mechanism by which sigma factors “time share” RNA polymerase complexes previously described in *B. subtilis* (*47*) might also be present in *E. coli*. Additionally, our results suggest that the pulses of activation of different sigma factors might be associated with different cellular growth statuses.

In future experiments, it would be interesting to conduct measurements using pairs of reporters so that expression pulses can be monitored simultaneously in the same cell. This is particularly interesting to examine in relation to the growth rate, as we observed many instances of anti-correlation, but the temporal properties of these pulses were distinct from each other. Although we measure growth rate in our analysis, our findings do not preclude the possibility that growth rate is a secondary effect that is downstream of the true coordinating signal, such as metabolic state (*48*). Experiments that use reporters to measure genes involved in metabolism alongside the stress response reporters or studies that carefully control the nutrient composition could be used to investigate this possibility. It will also be interesting to investigate how the information encoded in transcription factor dynamics is transmitted to downstream genes, and emerging technologies that enable the manipulation of gene expression over time could help to identify how these signals are propagated (*49, 50*). Finally, although we have focused our analysis of stress survival on transient ciprofloxacin treatment in wild type cells with the *gadX* reporter, it would be interesting to test survival to other antibiotics, different stress application patterns, and the effect of different strain backgrounds.

In summary, our findings reveal that pulsatile dynamics in gene expression and growth serve as a mechanism that cells can leverage to transiently resist stress. This dynamic behavior is spread across a broad range of stress response genes, including heat-shock response, multi-drug resistance, and oxidative stress and can enable subpopulations of cells to withstand temporary stress.

## Materials and Methods

### Strains and growth media

Strains containing reporter plasmids were sourced from the collection created by Zaslaver *et al*. (*51*), unless otherwise noted. Briefly, each strain has a low copy number plasmid (SC101 origin) containing the promoter sequence for the gene of interest upstream of the coding sequence for *gfpmut2* green fluorescent protein (GFP). The strain reporting *purA* was sourced from Rossi *et al*. (*7*) and the strain reporting *marA* was sourced from El Meouche *et al*. (*6*), where the *marA* reporter has intact MarR binding sites. The *purA* and *marA* reporters use a low copy number plasmid (SC101 origin) and the promoters of interest control the expression of cerulean cyan fluorescent protein (CFP). The constitutive reporter plasmid was constructed by cloning a constitutive promoter from Ref. (*52*) into the empty vector from the Zaslaver *et al*. library, pUA66. Specifically, the promoter used here is a variant of the σ70 consensus sequence: TTATCAAAAAGAGTA<u>TTGTCT</u>TAAAGTCTAACCTATAG<u>GAAAAT</u>TACAGCC**A**TCGAG AGGGACACGGCGAA where the +1 transcriptional start site is shown in bold and the -10 and - 35 regions are underlined. In all cases, we used *E. coli* MG1655 as the strain background.

The strain containing an integrated *gadX* reporter was constructed as described by Patange *et al*. (*8*). A cassette containing the *gadX* reporter and a kanamycin resistance gene was amplified from the reporter plasmid using the primers 5’-ATAAACACGTTCGTGTCCCGACAGGCACACAGACGGTTAGCCACTAATTAGAGCTCTCGAACCCCAGAGT-3’ and 5’-GTAAGAATAAAAAAAACGGGTCACCTTCTGGCGACCCGTTTTTCTTTGCGCCTGCAGGTCTGGACATTTA-3’ and integrated into the chromosome between the *nupG* and *speC* genes.

The Δ*rpoS* strain was constructed as described by Baba *et al*. (*53*). Briefly, an in-frame deletion of *rpoS* was achieved through integration of a cassette containing a kanamycin resistance gene between FLP recognition target sites and 50 base-pair homology sequences for the regions flanking *rpoS:* 5’*-*TGAGACTGGCCTTTCTGACAGATGCTTACTTACTCGCGGAACAGCGCTTC-3’ and 5’-CTTTTGCTTGAATGTTCCGTCAAGGGATCACGGGTAGGAGCCACCTTATG-3’ After successful integration, the kanamycin resistance gene was excised using FLP recombinase.

Cells were grown in M9 medium (0.1 mM CaCl_2_; 2 mM MgSO_4_; 1X M9 salts) supplemented with 0.4% glucose, 0.2% casamino acids, and 30 μg/ml kanamycin for plasmid maintenance. Media used in microfluidic experiments were supplemented with 2 g/L F-127 Pluronic to prevent cell adhesion and growth outside of the mother machine growth chambers.

### Static fluorescence microscopy snapshots

Overnight cultures were diluted 1:100 and incubated at 37C with shaking for 2.5 to 4 hours (OD_600nm_= 0.7-1.3). 1 μL of cells were placed on 1.5% MGC (0.2% glycerol, 0.01% casamino acids, 0.15 μg/ml biotin, and 1.5 μM thiamine) low melting point agarose pads and imaged at 100X on a Nikon Ti-E inverted fluorescence microscope. Three separate overnight cultures were sampled and imaged for each reporter strain.

### Mother machine microfluidic device

The mother machine microfluidic master mold used was described previously in Ref. (*26*). Briefly, the master mold chip has 8 independent main feed channels where growth media flows in and out. Each channel features 1,000 one-ended chambers (L x W x H = 25 μm x 1.3 μm x 1.1 μm) where the mother cells are trapped. We made the microfluidic devices by pouring a degassed 10:1 mixture of dimethyl-siloxane monomer and curing agent (Sylgard 184 Silicone Elastomer Kit, Dow Corning) onto the wafer, which was then cured overnight at 65C. Individual chips were separated from the mold and a 0.75 mm biopsy punch was used to create inlets and outlets for each flow channel before the chip was plasma bonded to a glass slide.

### Time-lapse microscopy movies

Overnight cultures were diluted 1:100 and incubated at 37C with shaking for ∼3 hours until mid-exponential phase (OD_600nm_= 1.0-1.3). Cells were concentrated by centrifugation (6,000x*g* for 2 min) and loaded into the microfluidic chip. Cells were seeded in the growth chambers by centrifugation at 6,000x*g* for 3 min. Media was supplied at a flow rate of 20 μL/min using a peristaltic pump. Cells were allowed to adapt to growth in the device for 2-4 hours before imaging. Time-lapse movies were acquired with a 100X oil objective on a Nikon Ti-E inverted fluorescence microscope equipped with a perfect focus system and a temperature control chamber that was set at 37C for the duration of the experiment. Images were acquired in phase contrast and epifluorescence illumination with a GFP or CFP filter every 5 min. The total number of cell lineages tracked for each reporter is listed in Table S2.

### Microscopy image processing and fluorescence and growth rate analysis

For static snapshots, cell images were segmented with the SuperSegger software (*54*). For mother machine experiments, we performed automated cell segmentation and lineage tracking using the deep learning-based software DeLTA (*26*). In both cases, reported single-cell fluorescence values are the average fluorescence intensities of all pixels belonging to each cell. Unless otherwise noted, for mother machine experiments only data for the ‘mother’ cell trapped at the top of the growth chambers was used in quantitative analysis. We smoothed the fluorescence data using a moving average filter with a window of 5 frames.

To estimate growth rates, we calculated the difference in cell length between adjacent frames normalized by the cell length, 1/L dL/dt. Specifically, when there is no cell division event between frames, this is calculated as (L_t_ – L_t-1_)/ (L_t-1_ Δt), where L_t_ is the length of the cell at time t, L_t-1_ is the length at the preceding frame, and Δt is the time between frames (5 min). In cases where a cell division event occurs, this calculation is modified to (L_t_ + D_t_ – L_t-1_)/ (L_t-1_ Δt), where D_t_ is the length of the daughter at time t. After calculating the growth rate, we smoothed the data using a moving average filter with a window of 5 frames.

Cell division times are calculated as the time between division events.

### Pulse identification

To allow comparison across all reporters, which have different expression levels and where imaging exposure times are different (Table S2), we normalized each time-series by its mean. Pulses of gene expression were identified using the built-in MATLAB function *findpeaks*, which identifies local maxima. We set the threshold for the minimum peak prominence at 0.5, unless otherwise noted. This value was determined empirically (Fig. S4).

### Autocorrelation and cross-correlation calculations

We calculated the autocorrelation of the fluorescence signal—R_fluo,fluo_(τ) where τ is the time shift—using the MATLAB function *xcov* with the unbiased normalization method. We then normalized the entire signal by the value of the autocorrelation at zero time lag, R_fluo,fluo_(0). The half life was calculated by taking the mean of all autocorrelation signals from individual cell lineages and identifying the time shift value where it crosses 0.5.

The cross-correlation between the fluorescence signal and the growth rate, R_fluo,gr_(τ), was calculated using the MATLAB function *xcov* with the unbiased normalization method. The signal was then normalized by the square root of the product of the autocorrelation at zero time shift of the fluorescence and growth rate signals: (R_fluo,fluo_(0) R_gr,gr_(0))^1/2^.

R_extreme_ is the value of R_fluo,gr_(τ) with the largest absolute value. τ_extreme_ is the time lag value associated with R_extreme_.

### Antibiotic survival assay

Mother machine experiments were initiated using the methods described above. During image acquisition, cells were provided with fresh medium for 14 hours, followed by 35 minutes of treatment with medium supplemented with 2 μg/ml ciprofloxacin, then fresh medium again for 16 hours. The outcomes of ciprofloxacin treatment for each lineage were manually scored as: ‘survived,’ ‘died,’ or ‘filamented.’ Cells were scored as ‘survived’ if cell division was observed in the chamber at the end of the full 16 hours after the second addition of fresh medium. Cells were scored as ‘died’ if growth permanently ceased in the chamber at any point after antibiotic exposure. ‘Filamented’ cells were excluded from the analysis, as it was difficult to accurately assess their outcome since they were frequently swept out of the chamber and field of view.

## Acknowledgments

We thank members of the Dunlop Lab and Wilson Wong for helpful discussions; Ahmad (Mo) Khalil and Razan Alnahhas provided input on the manuscript. Razan Alnahhas and Eric South conducted initial experiments with the constitutive reporters.

## Author contributions

NMVS and MJD conceptualized the study. NMVS and CMB performed experiments. NMVS, CMB, JBL, and MJD performed data analysis. NMVS and MJD wrote the manuscript, with input from CMB and JBL. MJD supervised the study and acquired funding.

## Supplementary Figures

**Figure S1.**
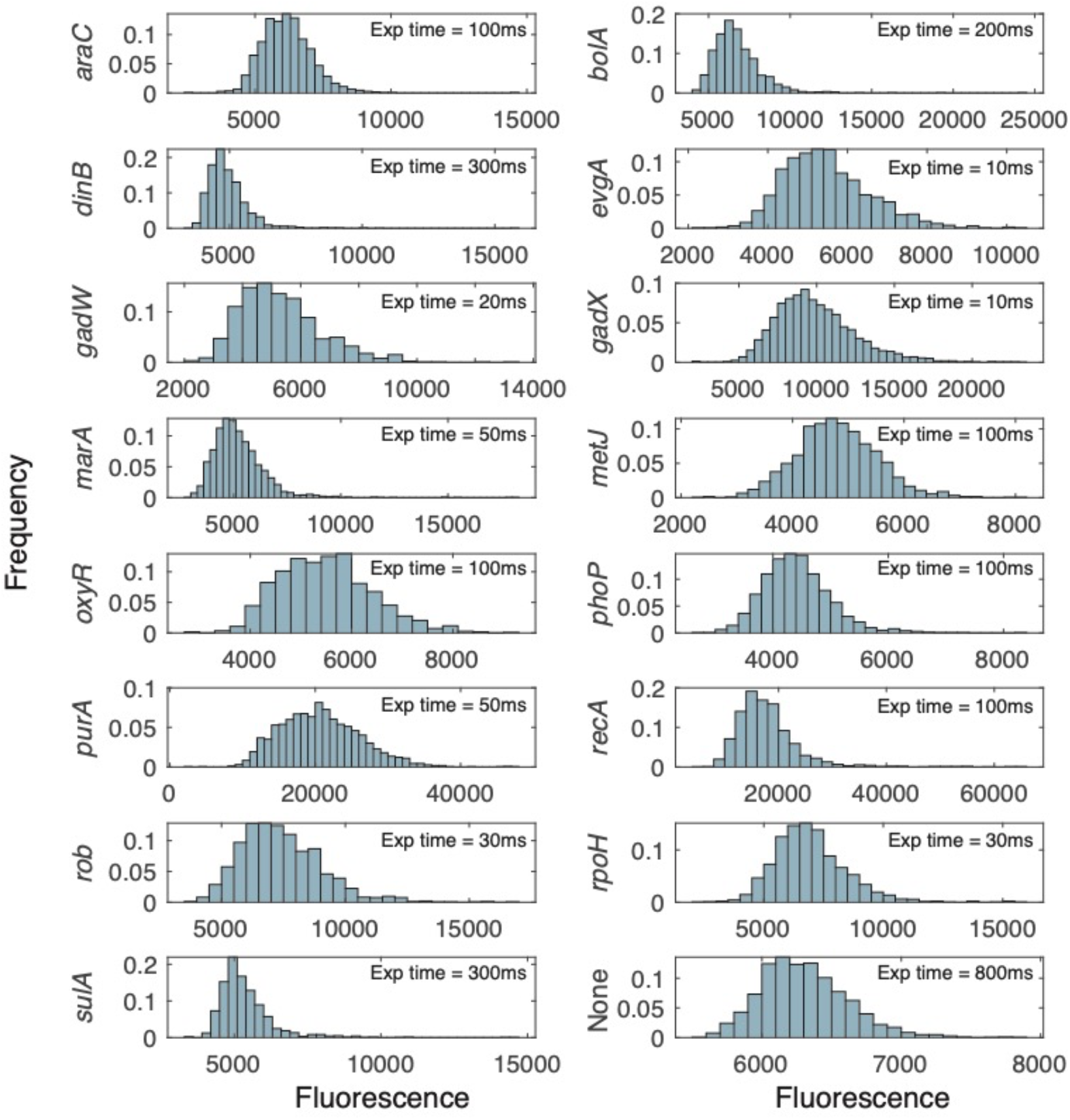
Histograms of fluorescence distributions of single cells. Microscopy exposure times vary between reporters to optimize detection of fluorescence from each reporter and are indicated in the figure.

**Figure S2.**
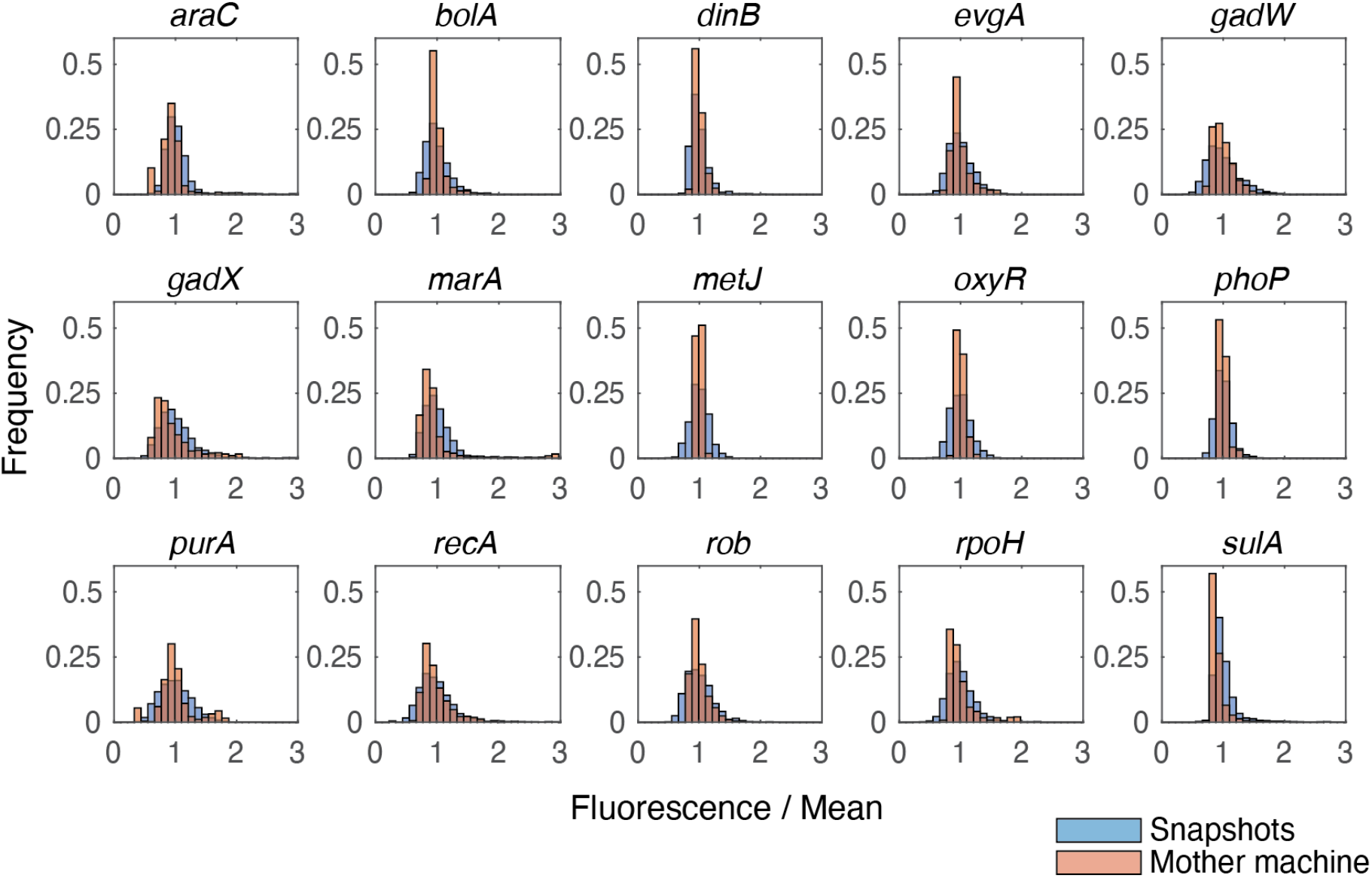
Histograms displaying distributions of mean normalized fluorescence values for single cells imaged in snapshots of agarose pads (blue) or in the mother machine device (red).

**Figure S3.**
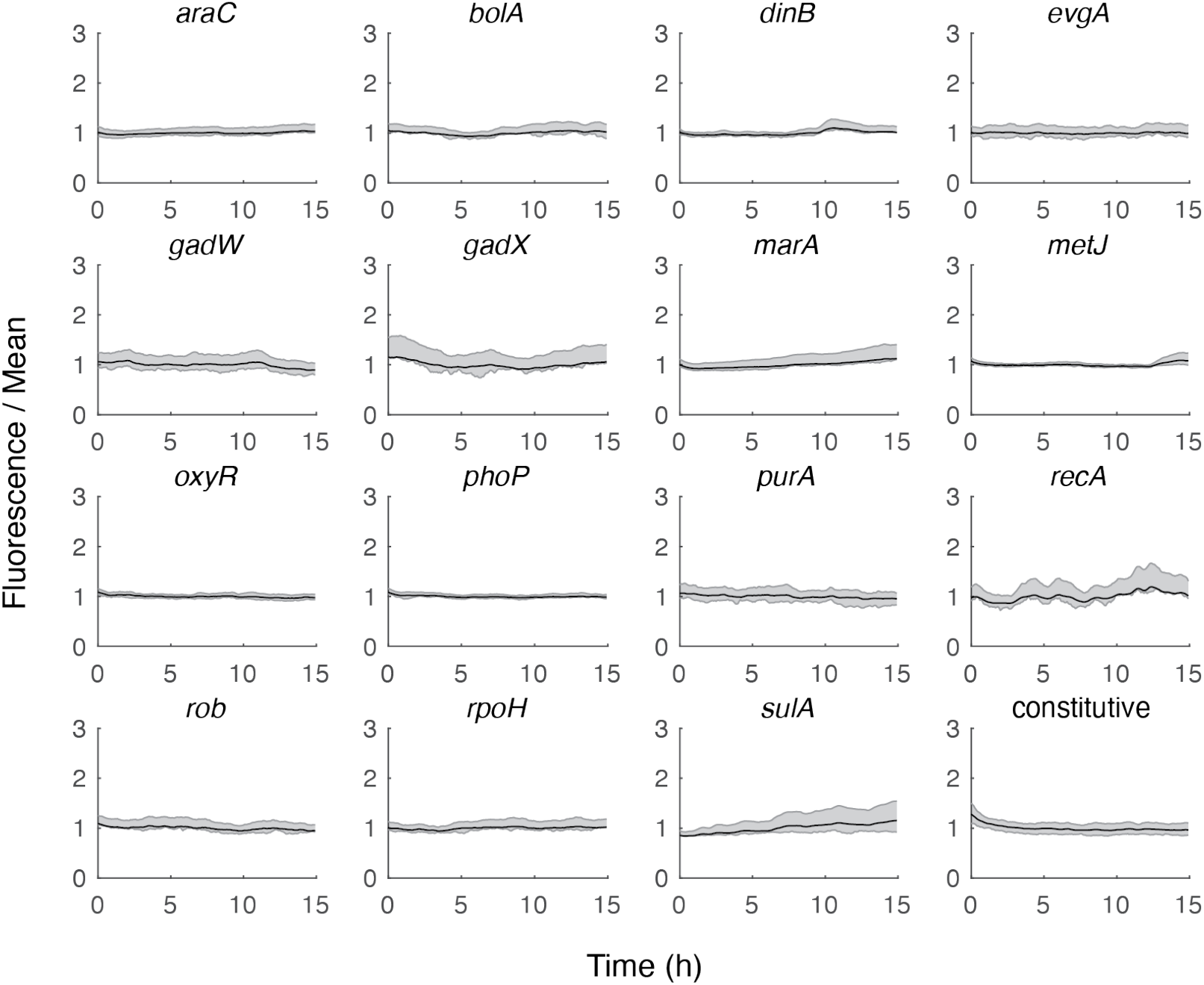
Mean fluorescence of all mother cells over time. Shaded area represents the 25 to 75 percentile values for each reporter; center line is the mean.

**Figure S4.**
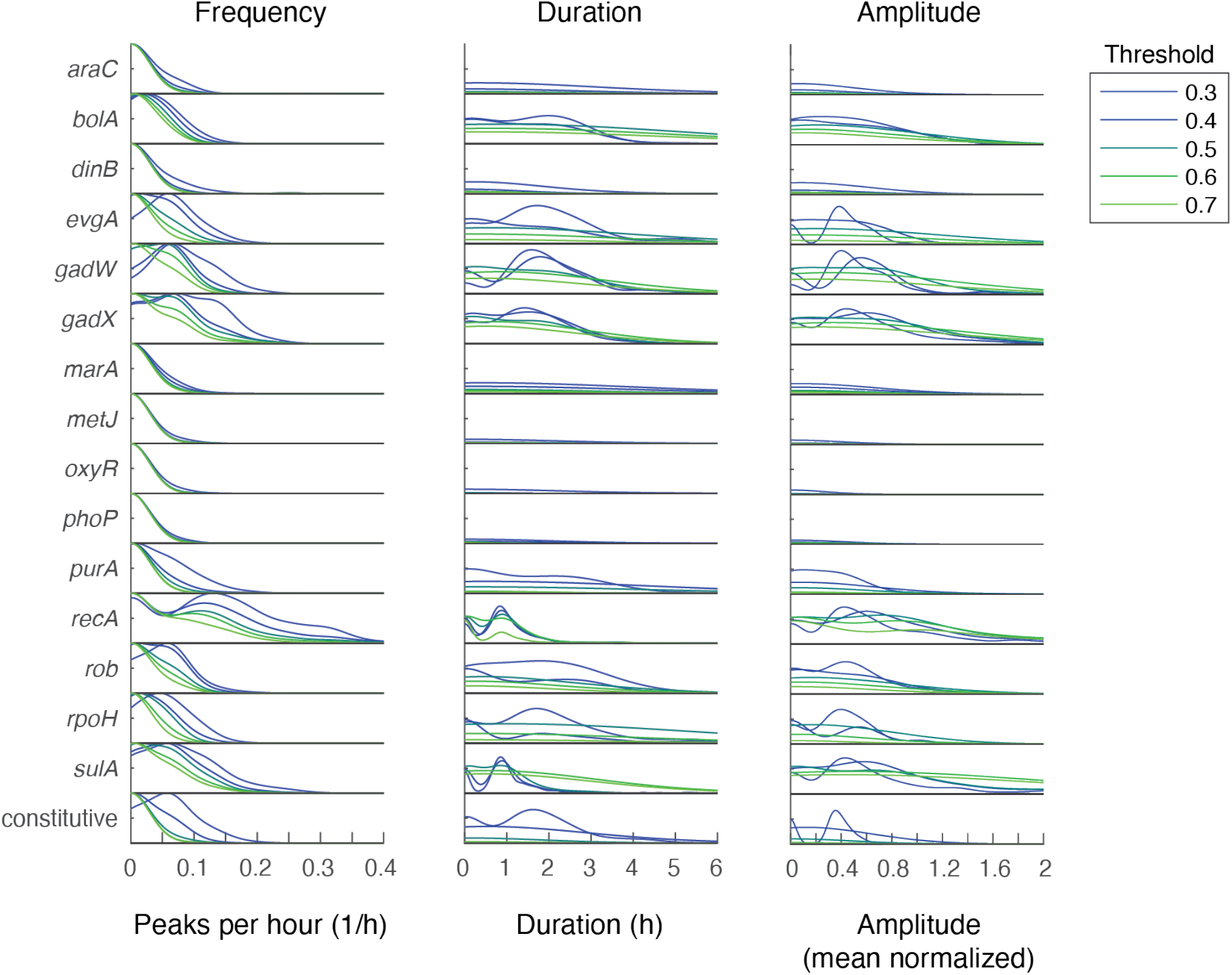
Effect of threshold value on pulse frequency, duration, and amplitude. Note that as the threshold is increased, we exclude small pulses that result from noise in the signal. This results in decreases in the frequency and generally increases in the duration and amplitude of the pulses. The default threshold value used for analysis is 0.5.

**Figure S5.**
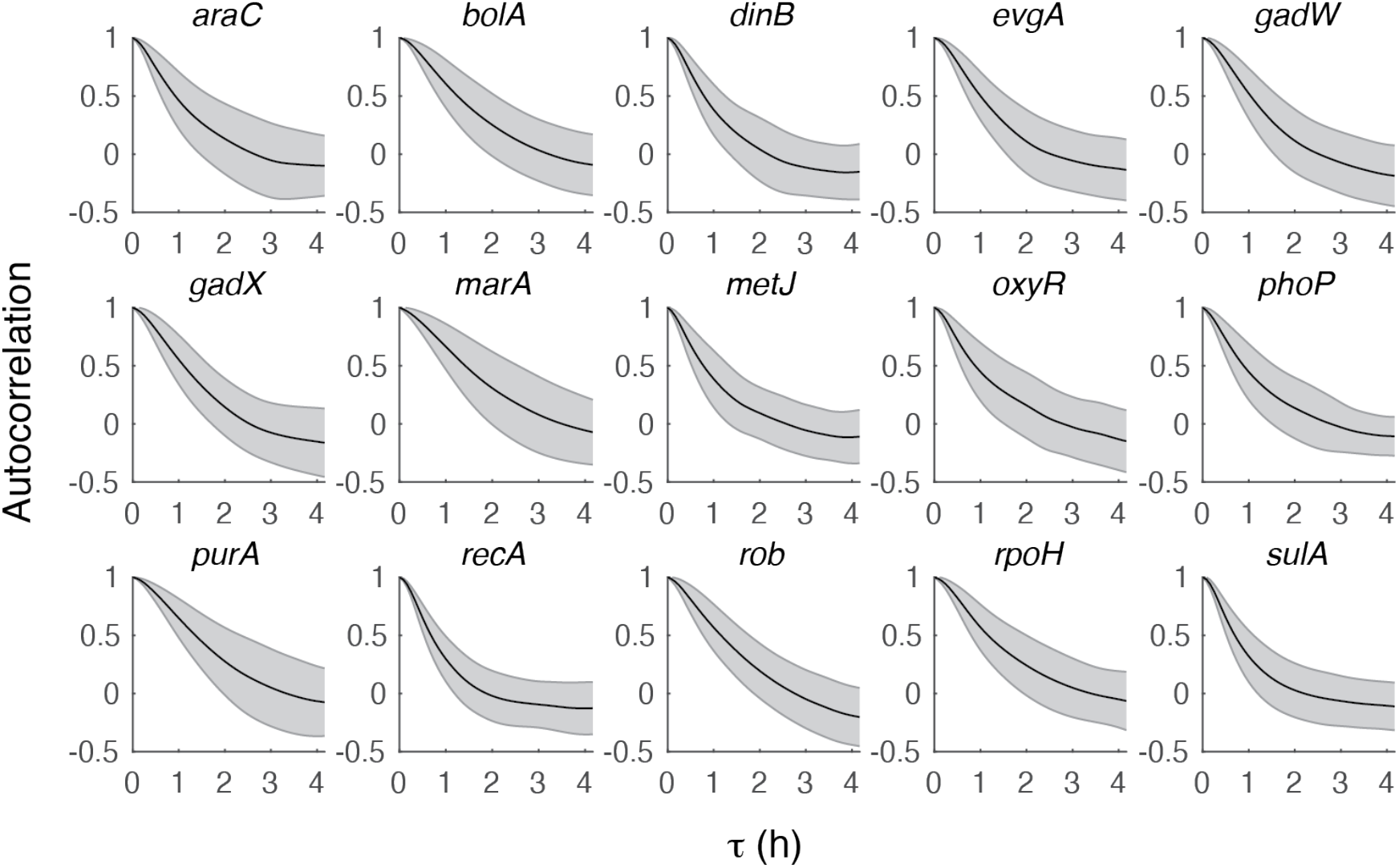
Autocorrelation of the fluorescence signal for all reporters. Shaded area shows standard deviation around the mean.

**Figure S6.**
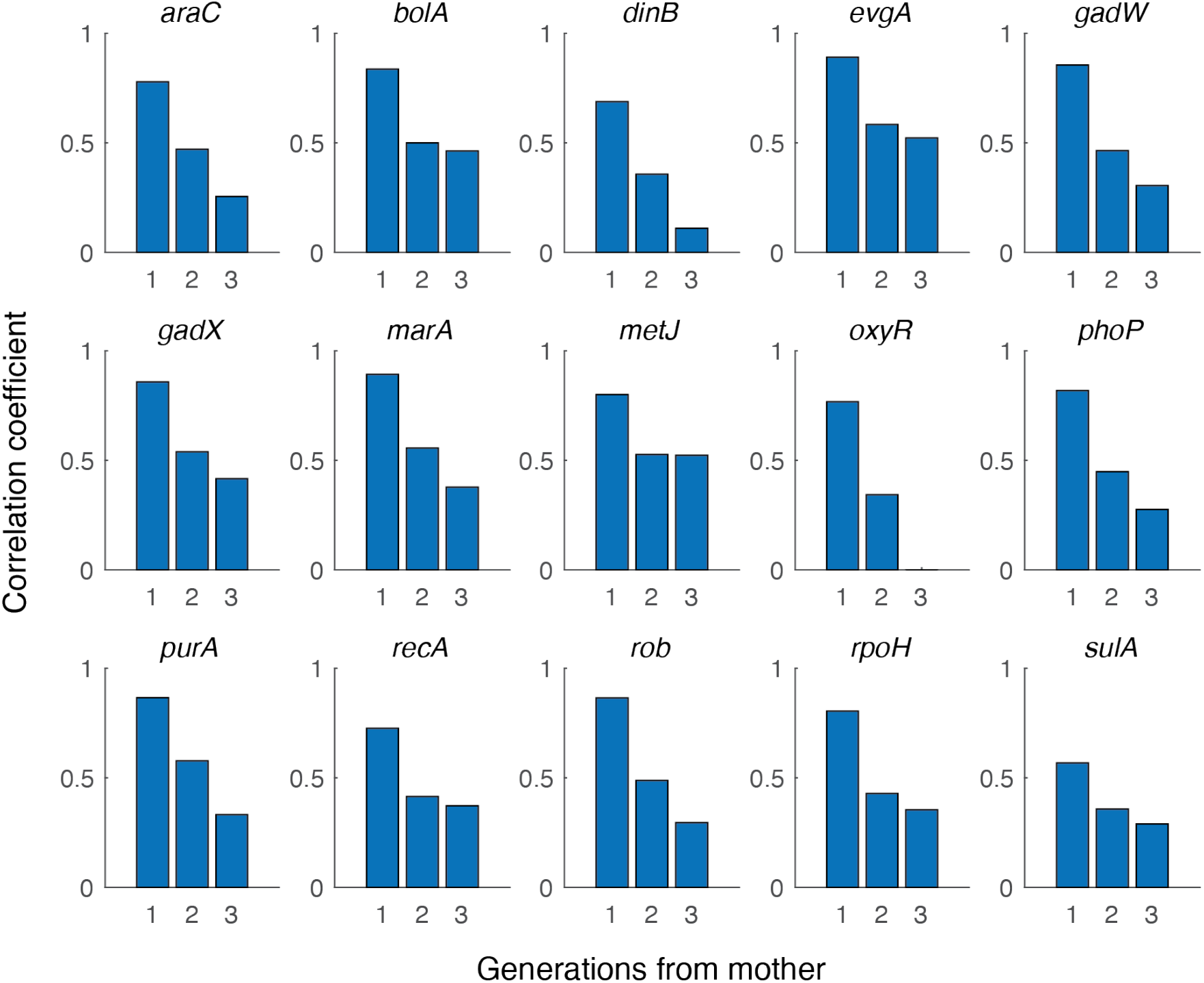
Correlation coefficient between mother cell’s mean fluorescence signal and its offspring. “Generations from mother” = 1 corresponds to a mother-daughter comparison, 2 is a mother-granddaughter comparison, and 3 is a mother-great granddaughter comparison.

**Figure S7.**
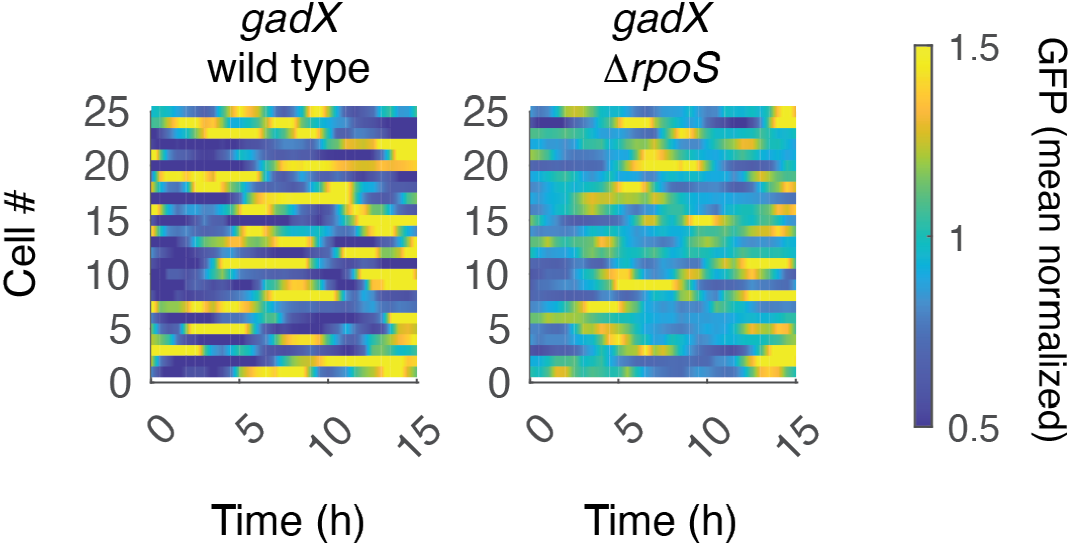
Single-cell measurements of green fluorescent protein expression (GFP) over time for the *gadX* reporter shown normalized relative to their mean in the wild type and Δ*rpoS* backgrounds.

**Figure S8.**
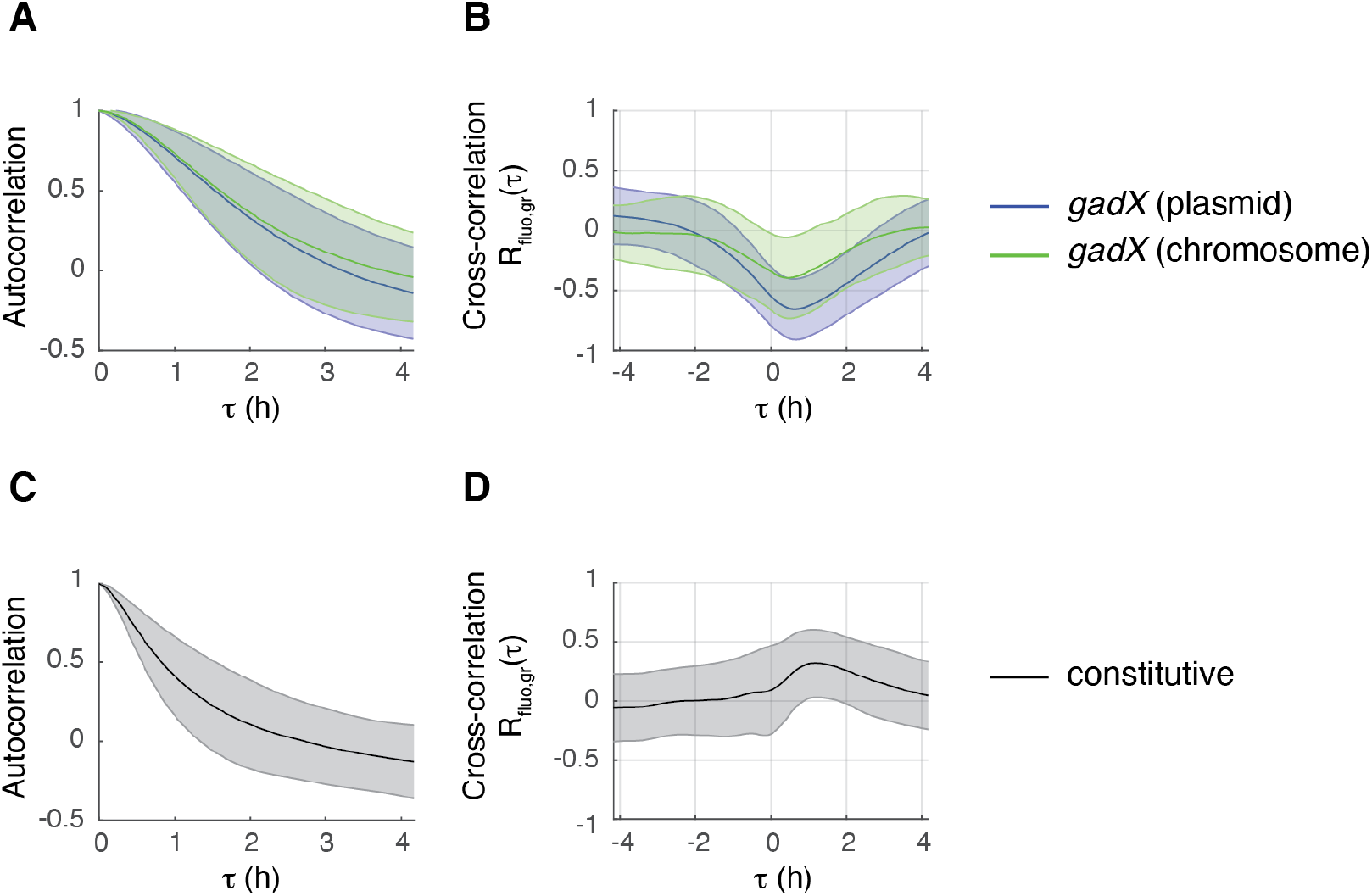
**(A)** Autocorrelation and **(B)** cross-correlation for the *gadX* reporter comparing plasmid-based and chromosomally-integrated versions. **(C)** Autocorrelation and **(D)** cross-correlation for the constitutive reporter. Shaded regions show standard deviation about the mean.

**Figure S9.**
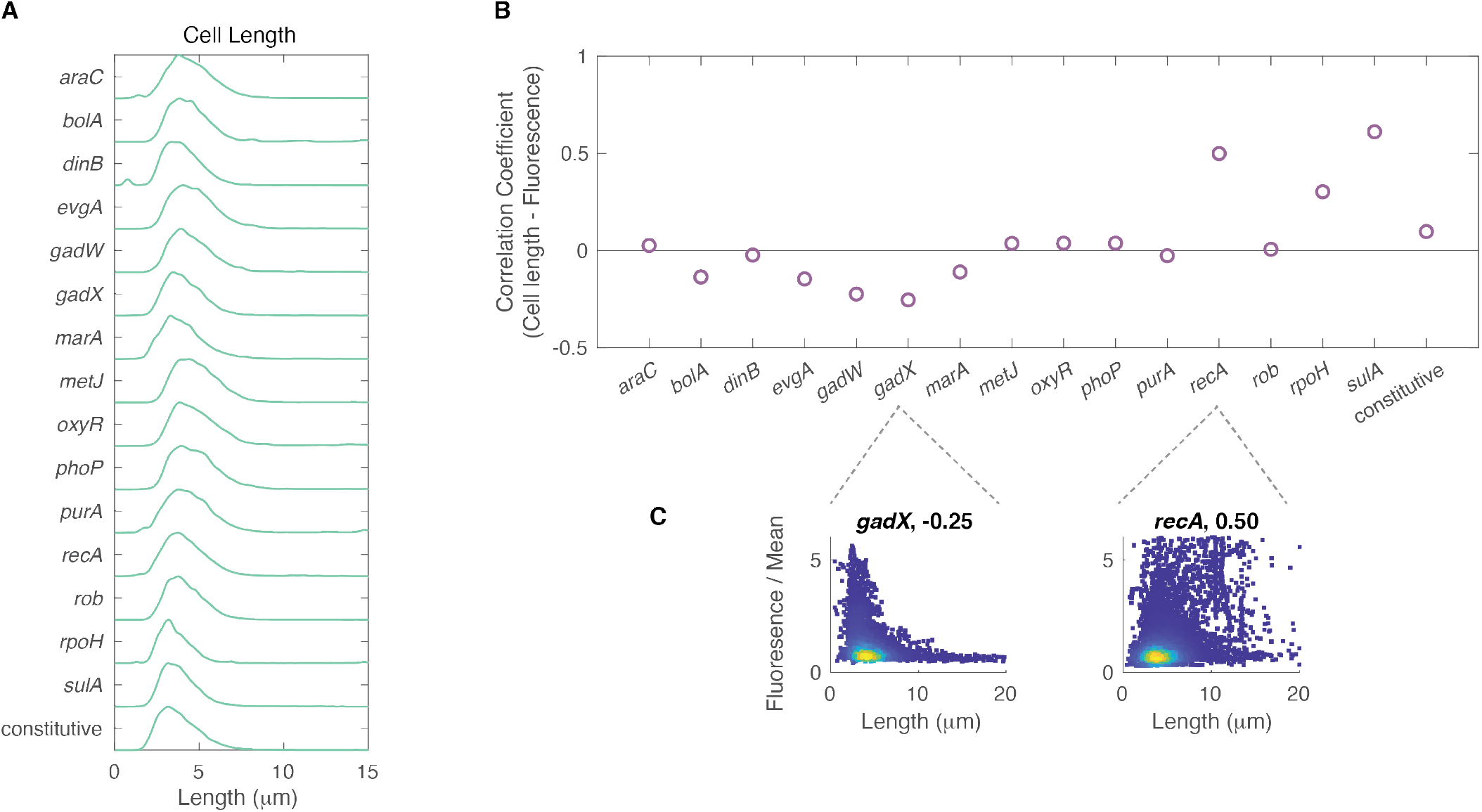
Relationship between cell length and fluorescence across reporters. **(A)** Distributions of cell length for all reporters. **(B)** Correlation coefficient values for cell length and fluorescence for all reporters. **(C)** As representative examples, insets show *gadX* or *recA* reporter levels plotted as a function of cell length for all time points and replicates. Correlation coefficient is listed in the figure title.

## Supplementary Tables

**Table S1.** Summary of genes characterized in this study

| Gene | Sigma Factor | Function | # of Regulatees |
| --- | --- | --- | --- |
| *araC* | RpoD | Arabinose catabolism | 14 |
| *bolA* | RpoD, RpoS | Cell morphology and protection, general stress response | 5 |
| *dinB* | RpoD, RpoS | SOS response, encodes DNA polymerase IV | 0 |
| *evgA* | RpoD, RpoS | Multi-drug resistance | 19 |
| *gadW* | RpoS | Acid resistance regulon | 15 |
| *gadX* | RpoD, RpoS | Acid resistance regulon | 34 |
| *marA* | RpoD | Multiple antibiotic resistance | 45 |
| *metJ* | RpoD | Methionine biosynthesis | 15 |
| *oxyR* | RpoD, RpoS | Oxidative stress | 42 |
| *phoP* | RpoH | Adaptation to low Mg<sup>2+</sup>, control of acid resistance genes | 56 |
| *purA* | RpoD | *de novo* synthesis of AMP | 0 |
| *recA* | RpoD | DNA recombination / repair protein, SOS response. | 0 |
| *rob* | Unknown | Resistance to antibiotics, organic solvents, heavy metals | 26 |
| *rpoH* | RpoD, RpoN, RpoE, RpoS | Sigma32. Response to heat shock during log-phase growth | 174 |
| *sulA* | RpoD | SOS response cell division inhibitor | 0 |

**Table S2.** Number of cell lineages tracked for each reporter, where each lineage corresponds to a single mother cell. Movie duration is the length of the measurement. Exposure time for the mother machine time-lapse experiments, which is customized for each reporter to account for differences in expression levels.

| Gene | # cell lineages tracked | Movie duration (median, in hours) | Exposure time (msec) |
| --- | --- | --- | --- |
| *araC* | 71 | 15.5 | 150 |
| *bolA* | 72 | 22.0 | 300 |
| *dinB* | 88 | 16.2 | 300 |
| *evgA* | 72 | 16.9 | 30 |
| *gadW* | 107 | 15.9 | 10 |
| *gadX* | 142 | 15.0 | 30 |
| *marA* | 450 | 16.3 | 50 |
| *metJ* | 71 | 15.5 | 150 |
| *oxyR* | 71 | 15.5 | 150 |
| *phoP* | 72 | 15.5 | 150 |
| *purA* | 101 | 16.8 | 50 |
| *recA* | 153 | 16.2 | 300 |
| *rob* | 108 | 15.9 | 10 |
| *rpoH* | 72 | 19.9 | 30 |
| *sulA* | 336 | 17.2 | 300 |
| constitutive | 216 | 16.3 | 350 |
| *gadX* chromosomal | 200 | 4.4 | 200 |
| *gadX* (WT) | 346 | 9.0 | 50 |
| *gadX* ($\Delta rpoS$) | 221 | 9.0 | 50 |

## Supplementary Movie Captions

**Movie S1**. Time-lapse movie of a representative mother machine chamber containing cells with the *gadX* reporter. Phase contrast (gray) and fluorescence channel (green) are shown. Time is displayed as HH:MM. Scale bar, 2 μm.

**Movie S2**. Time-lapse movie of a representative mother machine chamber containing cells with the *recA* reporter. Phase contrast (gray) and fluorescence channel (green) are shown. Time is displayed as HH:MM. Scale bar, 2 μm.

**Movie S3**. Time-lapse movie of a representative mother machine chamber containing cells with the *araC* reporter. Phase contrast (gray) and fluorescence channel (green) are shown. Time is displayed as HH:MM. Scale bar, 2 μm.

**Movie S4**. Surviving cell in a time-lapse movie where cells containing the *gadX* reporter are exposed to ciprofloxacin (period of treatment indicated with red bar at top of the image). Phase contrast (gray) and fluorescence channel (green) are shown. Time is displayed as HH:MM. Scale bar, 2 μm.

**Movie S5**. Dying cell in a time-lapse movie where cells containing the *gadX* reporter are exposed to ciprofloxacin (period of treatment indicated with red bar at top of the image). Phase contrast (gray) and fluorescence channel (green) are shown. Time is displayed as HH:MM. Scale bar, 2 μm.

